# Therapy-induced PSMA2 Sensitizes Prostate Cancer Cells to Residual Androgen and Promotes Neuroendocrine Lineage Transformation

**DOI:** 10.1101/2025.11.25.690552

**Authors:** N Patterson, A Badoi, G.P Vadla, M Hasani, J Moyer, C De la Nuez Ramirez, Y.C Chabu

## Abstract

Aberrant activation of the androgen receptor (AR) pathway drives prostate cancer (PCa). Androgen deprivation therapy (ADT) and next-generation AR blockade (e.g., enzalutamide) are initially effective, but virtually all patients develop castration-resistant prostate cancer (CRPC), which frequently transitions to treatment-emergent neuroendocrine PCa (tNEPC) following AR suppression. The molecular logic that links AR blockade to lineage plasticity remains incompletely understood.

Here, we identify PSMA2 (Proteasome Subunit Alpha 2) as a treatment-induced effector that mechanistically connects AR blockade to tNEPC evolution. Enzalutamide induces PSMA2 expression in AR-expressing PCa cells. Enforced PSMA2 expression accelerates HSP90 turnover, hypersensitizes AR to residual post-castration androgen, drives AR nuclear activity under androgen-poor conditions, and confers enzalutamide resistance. Conversely, PSMA2 silencing stabilizes HSP90, desensitizes CRPC to androgen, and re-sensitizes resistant cells to enzalutamide-induced cell death. Importantly, PSMA2 also promotes lineage plasticity: treatment-induced PSMA2 enhances transcriptional and phenotypic conversion toward tNEPC.

Thus, we uncover a single stress-induced node (PSMA2) that both maintains AR-dependent survival under ADT and fuels the neuroendocrine transition. PSMA2 marks an AR-hypersensitized transitional state and is itself a therapeutically actionable driver of tNEPC evolution, revealing an opportunity for rational interception of the lethal ADT-CRPC-tNEPC trajectory.

## INTRODUCTION

Androgen deprivation therapy (ADT) is foundational for prostate cancer management, yet nearly all patients progress to castration-resistant prostate cancer (CRPC). Emerging clinical, genomic, and transcriptional data support a therapy-induced lineage plasticity continuum in CRPC, in which potent ADT/AR pathway inhibition enriches for AR-low or canonical AR-indifferent transitional states that can further evolve into full treatment-emergent neuroendocrine prostate cancer (tNEPC) in a subset of patients^1–9^. Defining the molecular machinery that allows these transitional AR-low states to simultaneously sustain AR output under castrate androgen and acquire neuroendocrine identity remains a critical unmet need^9^.

The heat shock protein HSP90 plays an essential role in controlling ligand-dependent AR transcriptional activity: in the absence of androgen, HSP90 sequesters mature AR in the cytoplasm; with androgen present, AR disengages HSP90, translocates to the nucleus, and activates survival and growth gene programs^10,11^. Stabilization of the AR-HSP90 protein complex inhibits AR nuclear activity^12–15^. Enzalutamide is a potent AR antagonist that disrupts androgen-AR signaling and prolongs survival in CRPC, but it also imposes strong selective pressure that enriches for AR^low/−^, lineage-plastic clones capable of transdifferentiating into treatment-emergent neuroendocrine prostate cancer (tNEPC)^8,16–18^. This adaptive process is strongly associated with combined TP53/RB1 loss, N-MYC-driven epigenetic reprogramming, SOX2-/SPINK1-dependent plasticity programs, and expression of neuroendocrine genes (e.g.,*CHROMOGRANIN-A/CGA* and *SYNAPTOPHYSIN/SYP)* ^17,19–24^.

Here we identify proteasome subunit α2 (PSMA2) as a treatment-induced effector that links AR blockade to tNEPC evolution. PSMA2 is a core component of the 20S proteasome^25–30^, and is elevated in multiple cancers ^31,32^, yet its role in therapy adaptation is unclear. Our data support a model in which enzalutamide-induced PSMA2 alters proteasome subtype composition, enabling selective turnover of HSP90, thereby permitting AR nuclear signaling despite residual androgen. In parallel, PSMA2 drives stemness and neuroendocrine transcriptional programs, indicating that a single therapy-induced proteasome subunit coordinates both biochemical and fate-level axes of ADT resistance. This work identifies PSMA2 as a mechanistic driver of the AR-low to tNEPC conversion funnel and nominates PSMA2 as a candidate biomarker and therapeutically actionable node for rational interception of tNEPC evolution upstream of AR-indifferent disease.

## RESULTS

### Prostate cancer cells upregulate PSMA2 in response to enzalutamide

PSMA2 is overexpressed in multiple cancers^31,32^. We asked whether enzalutamide directly induces PSMA2 in prostate adenocarcinoma (LNCaP), AR-low transitional cells (LASCPC-01), and bona fide NEPC (NCI-H660)^33–37^. Quantitative polymerase chain reaction (qPCR) analyses revealed that enzalutamide transcriptionally activates PSMA2 across all three disease states, with the strongest induction in LASCPC-01 and NCI-H660 (Fig. 1A). Consistent with PSMA2 transcriptional activation, enzalutamide elevated PSMA protein levels in LASCPC-01 cells and in the TRAMP (Transgenic Adenocarcinoma of Mouse Prostate) mouse model of NEPC^38–41^ (Fig. 1B, 1C). Further, Analysis of TCGA patient data confirmed PSMA2 upregulation in human tumors (Fig. 1D) and PSMA2 gene amplification correlated with poor survival (Fig. 4N). Therefore, enzalutamide induces PSMA2 in human and murine prostate cancer, particularly in the AR-low and NEPC compartments most associated with therapy-induced lineage plasticity. Taken together, the above findings suggested that PSMA2 is part of an adaptive molecular response that promotes PCa resistance to enzalutamide and progression to tNEPC.

**Figure 1.**
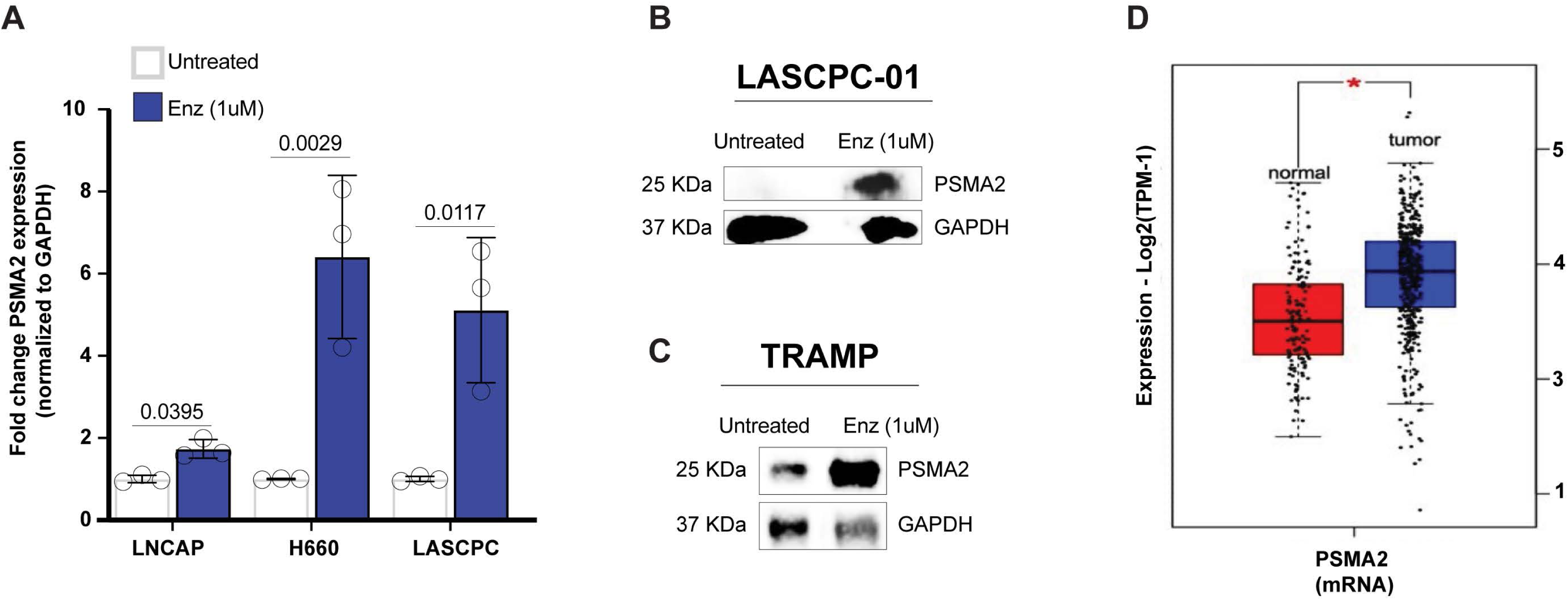
Prostate cancer cells upregulate PSMA2 in response to enzalutamide. (A) Quantitative PCR graph showing fold change expression of PSMA2 mRNA normalized to GAPDH in prostate cancer cells (LNCAP or LASCPC-01 or NCI-H660) treated with enzalutamide (1uM, 48 hours) (blue bars) versus untreated controls (grey bars). For this and all subsequent graphs p-values are derived from T-test (n.s=not significant). (B) Image of a Western blot showing PSMA2 abundance in LASCPC-01 cells left untreated or treated with enzalutamide (1uM) for 48hrs. Blots were stained against PSMA2 or GAPDH as loading control. (C) Western blot image showing expression of PSMA2 in TRAMP mouse prostate tumor tissue. Twenty-five-weeks-old B6FVB TRAMP mice received regular water (Control) or water containing enzalutamide (3mg/kg) (experimental) for 6 weeks. Prostates tissues were harvested and processed for protein extraction and Western blotting. Blots were stained against PSMA2 or β-Tubulin as loading control. (D) Box plot showing PSMA2 mRNA expression in TCGA and GTEx normal (red) versus tumor (blue) prostate tissues.

### Enzalutamide-resistant cells are sensitive to PSMA2 Inhibition

To directly test whether PSMA2 mediates enzalutamide resistance, we asked whether PSMA2 inhibition re-sensitizes prostate cancer cells to enzalutamide-induced cell death. We focused on LASCPC-01 cells because they are AR-low, resistant to enzalutamide, and exhibit adenocarcinoma–neuroendocrine lineage plasticity^34,42,43^. As expected, enzalutamide (1uM) had a negligible effect on the viability of intact LASCPC-01 cells in MTT (3-(4,5-dimethylthiazol-2-yl)-2,5-diphenyltetrazolium bromide) or Fluorescence-activated cell sorting (FACS)-based cell viability assays (Fig. 2A and 2E, respectively). In contrast, partial knockdown of PSMA2 using short hairpin RNA (PSMA2-KD) was sufficient to sensitize LASCPC-01 cell to enzalutamide-induced cell death (Fig. 2A, 2B). Note that moderate cell loss was observed following PSMA2-KD, however PSMA2-KD sensitized the surviving cells to enzalutamide-induced cell death (Fig. 2A).

**Figure 2.**
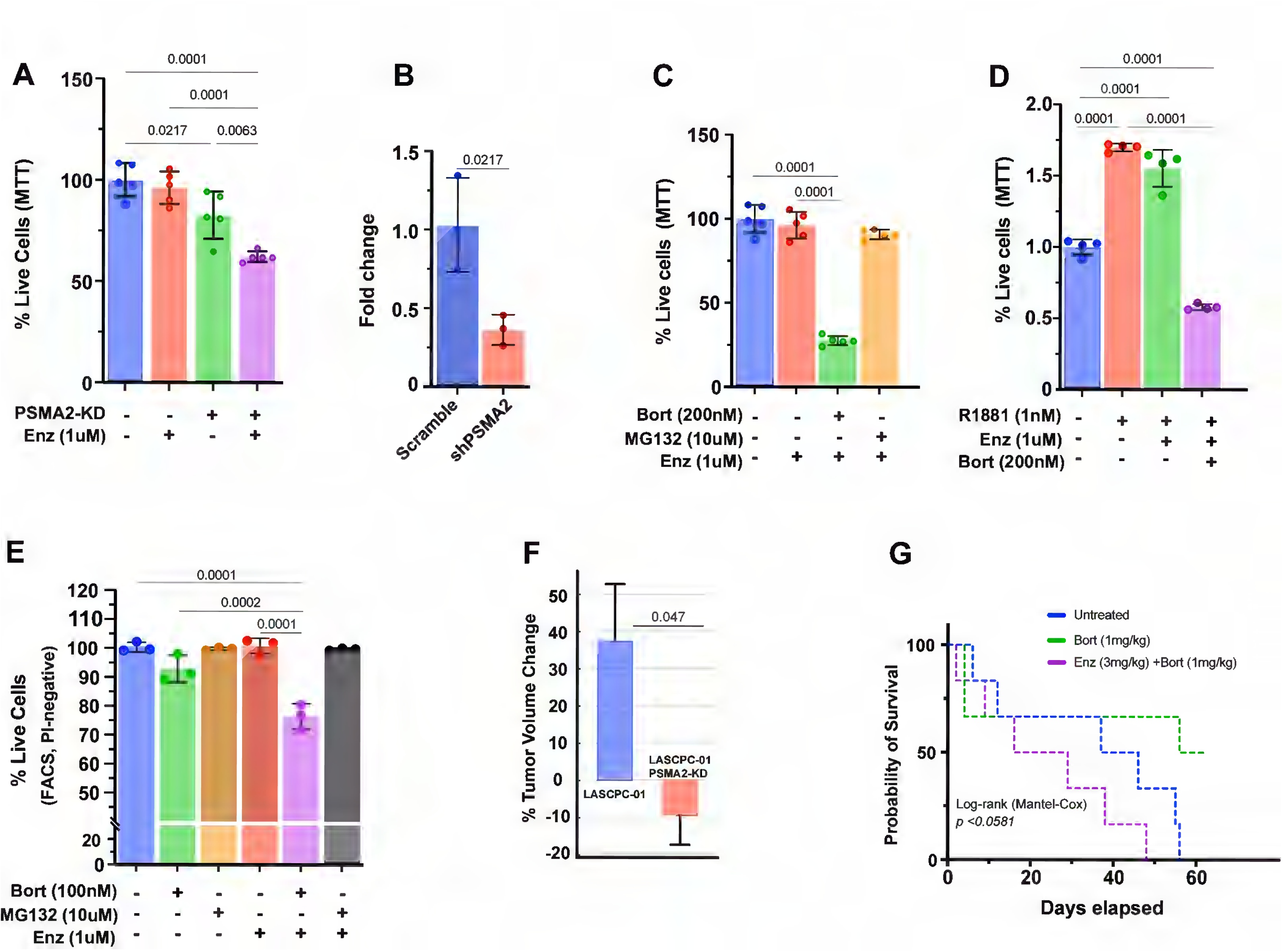
PSMA2 promotes ADT resistance. (A) MTT assays graph showing the relative viability of control LASCPC-01 cells or PSMA2-KD LASCPC-01 cells before or after enzalutamide treatments (1um, for 48hours) treatments (red). (B) Quantitative PCR graph showing fold change expression of PSMA2 mRNA normalized to GAPDH in LASCPC-01 transfected with scramble (control) versus PSMA2 short hairpin RNA. (C) MTT assay graph showing relative viability of LASCPC-01 cells left untreated or treated with enzalutamide (1uM for 48hours) alone or together with Bortezomib (200nm for 48hours) or MG132 (10uM for 3hours). (D) MTT assay graph showing relative viability of LASCPC-01 cells left untreated or treated with R1881 (1nM) in the presence of enzalutamide (1uM) alone or together with Bortezomib (200nm). (F) Graph showing the relative proportion of live LASCPC-01 cells in FACS experiments (%propidium iodide-negative cells). LASCPC-01 cells were treated with bortezomib or MG132 or enzalutamide single or combination treatments. (F) Flank tumors from nude Foxn^1u^ (nu/nu) mice bearing contralateral LASCPC-01 or LASCPC-01 PSMA2 KD flank tumors (N=4; 2 males + 2 females) derived from 10^5^ inoculated cells were measured once tumors reached ~15-90 mm^3^. Baseline volumes were recorded, and tumors were re-measured four days later to quantify percent change in tumor volume. Control tumors showed robust early expansion, whereas PSMA2-KD tumors displayed markedly blunted or negative growth, indicating a requirement for PSMA2 in early outgrowth. Bars represent mean SEM; significance was assessed by Welch’s t-test. (G) Kaplan Meier survivorship graph showing survival of 25-28 weeks-old B6FVB TRAMP mice left untreated or treated with Bortezomib alone or Bortezomib-enzalutamide combination treatments for two weeks.

PSMA2 is required for proteosome function ^28–30^. To assess whether the sensitization effect of PSMA2-KD is mediated by the proteosome, we asked to what extent pharmacological inhibition of the proteosome phenocopies PSMA2-KD. LASCPC-01 cells were treated with enzalutamide (1uM) in the absence or presence of the proteosome inhibitor Bortezomib (100 or 200uM). FACS cell viability assays showed that Bortezomib (200nM) killed enzalutamide resistant LASCPC-01 cells (Fig.2C). This effect is specific to Bortezomib as Carbobenzoxy-L-leucyl-L-leucyl-L-leucinal (MG132, 10uM), showed no detectable cell killing effect on enzalutamide resistant cells (Fig. 2C). These divergent effects are mechanistically informative as bortezomib and MG132 have distinct proteasome subtype specificity (see discussion).

Residual androgen was sufficient to drive the growth of LASCPC-01 cells even in the setting of enzalutamide. LASCPC-01 cell growth was evaluated in the presence of the synthetic androgen methyltrienolone (R1881, 1nM), Bortezomib (100nM), and enzalutamide (1uM). Bortezomib abrogated androgen mediated growth of enzalutamide resistant LASCPC-01 cell growth (Fig. 2D). Importantly, we detected a synergistic cell killing effect between Bortezomib, but not MG132, and enzalutamide (Fig. 2E).

Further, in vivo studies where LASCPC-01 cells are inoculated subcutaneously in the flank of nude (Foxn^1u^) mice showed that tumors lacking PSMA2 failed to expand in the androgen-limited microenvironment, whereas control tumors increased in volume (Fig.2F). This establishes PSMA2 as a critical determinant of early androgen-independent outgrowth, even prior to enzalutamide exposure.

We next evaluated how these early differences influenced disease outcome under enzalutamide pressure. Bortezomib treatment extended the survival of CRPC-NEPC TRAMP mice^38^ (Fig.2G). Interestingly, addition of enzalutamide blocked the life extension benefit of bortezomib, pointing to a toxic interaction between these two agents (see discussion).

### PSMA2 sensitizes CRPC to residual androgen by antagonizing the HSP90

Residual (post-castration) androgen continues to fuel CRPC growth via both canonical and non-canonical mechanisms^44–48^. We asked whether treatment-induced PSMA2 plays a role in this phenomenon. LASCPC-01 or PSMA2-KD LASCPC-01 cells were exposed to sub-nanomolar androgen (R1881) in the presence of enzalutamide. Parental LASCPC-01 cells proliferated rapidly in response to enzalutamide and increasing residual androgen levels, consistent with enzalutamide-induced PSMA2 (Fig. 3A). In contrast, PSMA2-KD cells were insensitive to androgen under enzalutamide (Fig. 3A). Thus, PSMA2 induction hypersensitizes AR to residual androgen.

**Figure 3.**
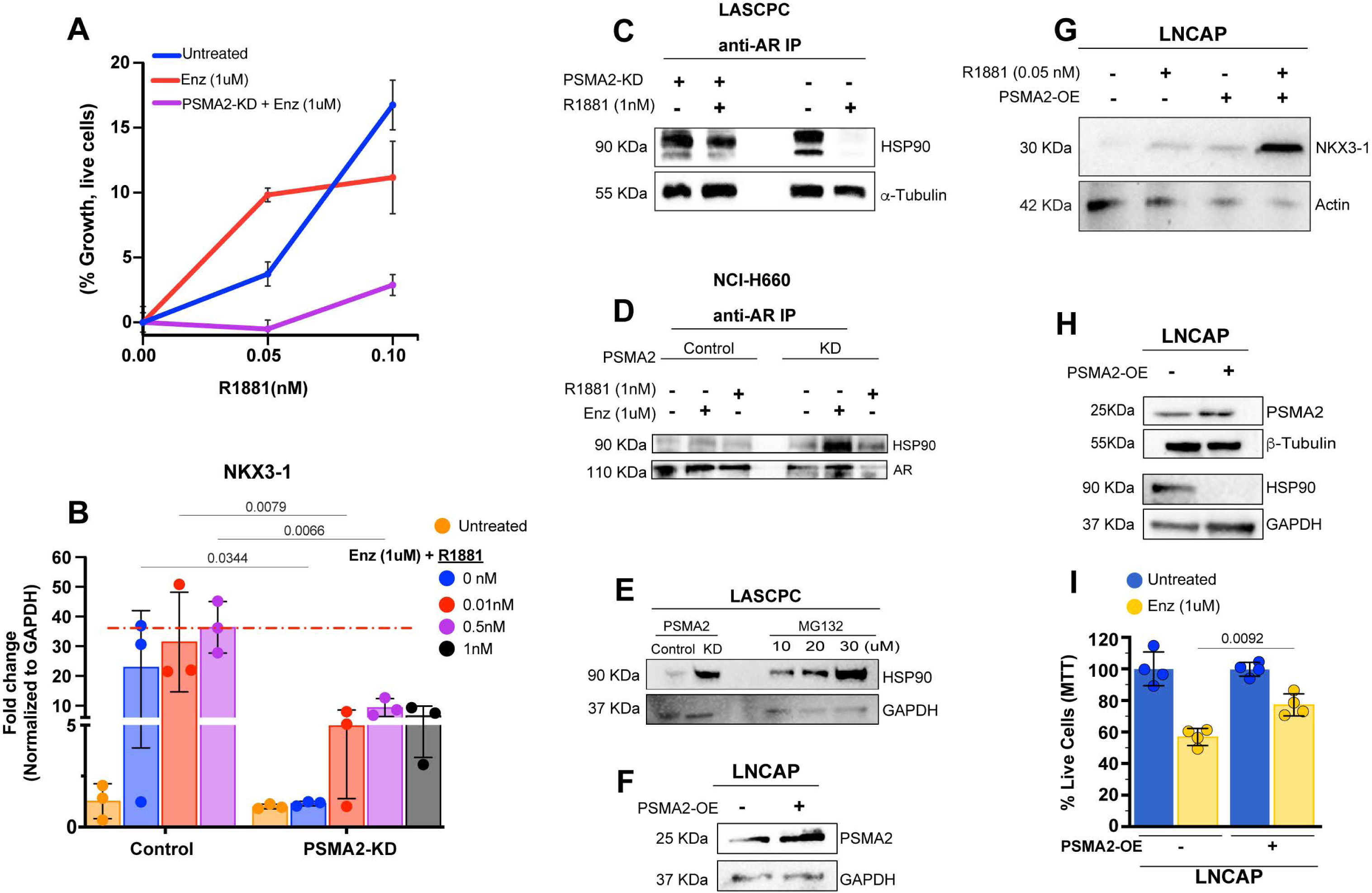
Treatment-induced PSMA2 antagonizes Hsp90-AR complex formation by promoting HSP90 turnover. (A) Trypan blue cell viability assay showing androgen growth response of LASCPC-01 or PSMA2-KD LASCPC-01 cells. Cells were treated with R1881 (0.05 or 0.1nM) in the absence (Untreated) or presence of enzalutamide (1uM). (B) Quantitative PCR graph showing *NXK3-A* expression in controls LASCPC-01 cells or PSMA2-KD LASCPC-01 cells treated with increasing levels of R1881 (0.01-1nM) in the presence of enzalutamide (1uM). Expression was normalized to GAPDH mRNA. (C and D) Western blot images from anti-AR immunoprecipitation (IP) assays on lysates derived from intact or PSMA2-depleted (PSMA2-KD) LASCPC-01 (c) or NCI-H660 (d) cells. Blots were stained with antibodies against Hsp90 or AR or β-tubulin. (E) Western blot image showing Hsp90 protein abundance in intact or PSMA2-depleted LASCPC-01 before and after treatments with increasing concentration of MG132 (10, 20, 30uM). Blots were stained with antibodies against Hsp90 or GAPDH (loading control). (F and G) Western blot image showing that direct partial over-expression of PSMA2 (F) is sufficient to sensitize LNCAP cells to androgen signaling-dependent upregulation of NKX3-1 (G). (H) Western blot images of lysates derived from controls or PSMA2-overxpressing LNCAP cells. Blots were stained against PSMA2 or or HSP90 or loading controls (β-tubulin and GAPDH, respectively). (I) MTT cell viability graph showing the relative viability of unaltered or PSMA2 over-expressing LNCAP cells following enzalutamide treatment.

We next examined AR transcriptional output using NKX3.1 expression, a dominant target in tNEPC^49^. LASCPC-01 markedly upregulated NXK3-1 in response to sub-nanomolar androgen (0.01-0.5 nM) Fig. 3B. In PSMA2-KD LASCPC-01 cells, NKX3.1 induction was sharply blunted, even when androgen concentrations were increased 100-fold (1 nM) (Fig. 3B; 0.01nM R1881, LASCPC-01 versus 1nM R1881 PSMA2-KD LASCPC-01). Therefore, PSMA2 is required for hypersensitive AR transcriptional responses to residual androgen.

HSP90 is a central regulator of AR activity: in the absence of ligand, HSP90 sequesters mature AR in the cytoplasm, preventing AR signaling^12–14^. We asked whether PSMA2 controls AR signaling by regulating the formation of AR-HSP90 protein complex. Protein extracts derived from controls or PSMA2-KD cells were used in anti-AR pulldown experiments. Western blotting determined the relative abundance of precipitated HSP90 protein across treatment conditions. As expected, we found that AR pulls down HSP90 but androgen treatment dramatically reduces the abundance of HSP90 protein co-precipitates (Fig. 3C, lane 3 vs. 4). However, in LASCPC-01 PSMA2-KD cell lysates, AR robustly pulls down HSP90 even following androgen treatment (Fig. 3C lanes 1 vs. 2 and compared to lanes 3 and 4). Similar results were obtained from the AR low NEPC cells NCI-H660 (Fig. 3D, compare HSP90/AR ratios). Thus, PSMA2 inhibition stabilizes the formation of an androgen-insensitive AR-HSP90 protein complex. Given PSMA2 known role in proteosome function and the observation that the proteasome-targeted agent Bortezomib mimics PSMA2-KD in term of re-sensitization to enzalutamide mediated cell death, we considered the possibility that PSMA2 normally destabilizes HSP90-AR protein complex formation by promoting HSP90 turnover. Indeed, we found that PSMA2-KD stimulates HSP90 protein abundance in LASCPC-01 cells (Fig. 3E). MG132-mediated proteosome inhibition increased HSP90 protein abundance in a dose-dependent manner (Fig. 3E, F). Conversely, moderate overexpression of PSMA2 was sufficient to stimulate androgen-dependent NKX3-1 upregulation (Fig. 3G). This effect was accompanied with dramatically reduced Hsp90 protein levels (Fig. 3H) and a corresponding desensitization of LNCAP cells to enzalutamide-induced cell death (~20%, Fig. 3I). Collectively, the above data argue that enzalutamide-induced PSMA2 promotes Hsp90 turnover to control androgen mediated CRPC growth.

### Treatment-induced PSMA2 acts as a lineage plasticity factor

CRPC progression under AR blockade is associated with acquisition of neuroendocrine fate^8,9,50,51^. We therefore asked whether treatment-induced PSMA2 contributes to lineage plasticity. Androgen blockade stimulates the Serine Peptidase Inhibitor, Kazal type 1 (SPINK1), which dominantly drives lineage plasticity and gives rise to NEPC ^23^. qPCR analyses showed that enzalutamide stimulated expression of SPINK1 and neuroendocrine genes (CGA and SYP) in LASCPC-01 cells. Importantly, this induction required PSMA2: PSMA2-KD abrogated enzalutamide-induced SPINK1, CGA, and SYP expression (Fig. 4A-C). Concordant with these results, enzalutamide elevated SYP protein levels in vivo in a PSMA2-dependent manner in immunohistochemistry from TRAMP mice (Fig. 4D-K). These findings indicate that PSMA2 is necessary for the lineage switching transcriptional program induced by enzalutamide.

**Figure 4.**
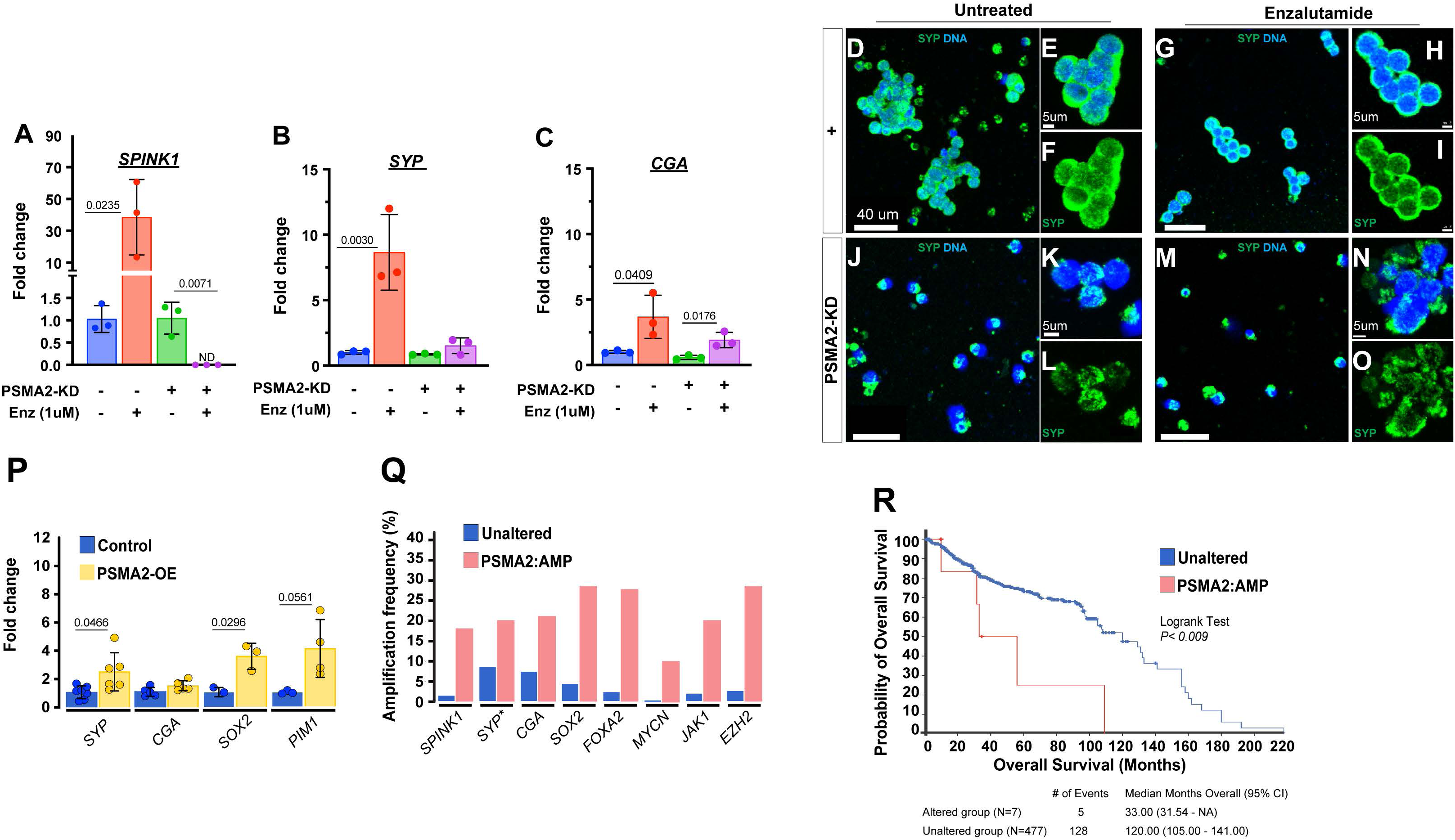
Treatment-induced PSMA2 contributes to NEPC transformation. (A-C) Quantitative PCR graphs showing the expression of *SPINK1* (A) or *SYP* (B) or *CGA* (C) in PSMA2 intact or PSMA2-depleted (PSMA2-KD) LASCPC-01 cells before and after enzalutamide treatments (1uM for 48hrs). All target mRNA expression levels were normalized to GAPDH. (D-O) Representative images of intact or PSMA2-depleted (PSMA2-KD) LASCPC-01 cells left untreated or treated with enzalutamide. Cells were fixed, co-stained against Synaptophysin (Syp) and DAPI (to detect nuclei), and imaged on a confocal microscope using identical imaging parameters. Low magnification images are presented in D,G,J, and M and corresponding higher magnification images are shown E,F, H, I versus K,L,N,O) (P) Quantitative PCR graph showing the relative expression of NEPC and stemness markers (*SYP*, *CGA*, *SOX2*, *PIM1*) in LNCAP cells over-expressing PSMA2 (yellow) compared to LNCAP controls (blue). All target mRNA expression levels were normalized to GAPDH mRNA. (Q) Graph showing the amplification frequency of NEPC and stemness genes in PSMA2 amplified versus PSMA2 unaltered prostate cancer patients using the cBioportal cancer genomics analysis platform. (R) Kaplan Meier survivorship analysis of PSMA2-amplified versus unaltered prostate cancer patients from cBioportal cancer genomic analysis.

We next asked whether PSMA2 is sufficient to drive lineage plasticity. Overexpression of PSMA2 in LNCaP cells stimulated expression of stemness and NEPC-associated genes including *SOX2, PIM1, SYP, CGA*, and *NSE* (Fig. 4L). Thus, PSMA2 is sufficient to induce neuroendocrine-fate transcriptional programs in an AR-dependent adenocarcinoma background.

Finally, analysis of patient genomic features using cBioPortal^52,53^ showed that PSMA2 amplification concords with NEPC transcriptional state and poor outcome (Fig. 4M-N). Collectively, these data support that treatment-induced PSMA2 is both necessary and sufficient to drive therapy-induced lineage plasticity toward NEPC.

## DISCUSSION

ADT drives prostate adenocarcinoma into an adaptive AR-low transitional state that frequently progresses to tNEPC, one of the most aggressive and therapeutically recalcitrant disease endpoints in prostate cancer^5,8,9^. This transition occurs despite extremely low androgen. It is unclear why AR transcriptional output persists in this window and how this same window enables lineage conversion.

Our data identify PSMA2 as a treatment-induced node that directly links these two processes. Enzalutamide induces PSMA2 expression in human and murine prostate cancer. PSMA2 induction enables residual-androgen hypersensitization by accelerating HSP90 turnover, releasing AR from cytoplasmic sequestration, and thereby enabling AR nuclear activation despite castrate androgen. Parallel to this, PSMA2 is necessary and sufficient to induce stemness and NEPC transcriptional programs, indicating that the same PSMA2 axis that sustains AR output under low androgen also opens the gate into the neuroendocrine lineage. We therefore define a PSMA2-driven signaling node that converts AR-low to NEPC state.

Mechanistically, although PSMA2 is a core proteasome subunit, the bortezomib vs MG132 divergence in our study suggests that enzalutamide-induced PSMA2 alters proteasome subtype composition to bias substrate specificity, selectively enhancing degradation of HSP90. PSMA2 overexpression promotes HSP90 turnover; PSMA2 loss or pharmacologic inhibition rescues HSP90 stability, re-imposes HSP90-AR sequestration, and extinguishes residual-androgen hypersensitivity.

Although MG132 stabilizes HSP90, confirming that HSP90 turnover is proteasome-dependent, it fails to phenocopy the bortezomib-enzalutamide synthetic lethality. This difference could be because Bortezomib is a slowly reversible inhibitor that durably targets the β5/β1 activities of the PSMA2-enriched proteasome subtype over 48 h, whereas MG132 activity is short lived at non-toxic dose^54–56^. Thus, transient global proteasome inhibition could be sufficient to reveal HSP90 as a PSMA2-proteasome substrate but insufficient to recapitulate the sustained proteasome subtype blockade and downstream stress responses required for cell killing.

We propose that PSMA2 acts early in the adenocarcinoma-AR^low^-NEPC trajectory, and several independent features of NEPC biology support this interpretation. Established tNEPC retains high HSP90 protein because N-Myc, AURKA, AKT, and other NE drivers are obligate HSP90 clients^22^. Also, clinical genomic datasets show enrichment of heat-shock and proteostasis programs in human NEPC ^5,8^. This means the terminal NEPC state requires abundant HSP90, not diminished HSP90. Consequently, the selective HSP90 turnover we observe with PSMA2 fits best within the transitional AR^low^ time frame, where modest reductions in AR-HSP90 complex stability are sufficient to free AR from cytoplasmic sequestration and enable residual-androgen hypersensitivity. Because proteostasis feedback may rapidly restores total HSP90, this early remodeling will not necessarily manifest as a detectable bulk HSP90 drop, yet it can still decisively alter lineage trajectory at the level of client flux and complex stability.

This early positioning is reinforced by the finding that PSMA2 overexpression alone is sufficient to induce NEPC-associated transcriptional programs (SOX2, PIM1, SYP, CGA) in an otherwise AR-dependent adenocarcinoma background.

This sufficiency indicates that PSMA2 does not simply mark established NEPC but can drive the lineage-permissive, chromatin-plastic intermediate state that precedes full neuroendocrine commitment. This aligns with prior work showing that proteostasis stress and proteasome remodeling can activate lineage-plastic programs in prostate cancer^57,58^, placing PSMA2 within a broader framework in which stress-responsive proteasome changes license NE-associated transcriptional states. Together, these data place PSMA2 at the mechanistic entry point of the AR^low^-NEPC conversion cascade and suggest that PSMA2 levels may serve as a quantitative biomarker of early lineage drift before irreversible NEPC fixation.

These findings have translational implications. First, PSMA2 induction could serve as an early molecular biomarker of the AR-low transitional state preceding tNEPC, which is the window in which most therapeutic opportunity still exists. Second, our data indicate that PSMA2-dependent proteasome specialization is therapeutically actionable: bortezomib was sufficient to extend survival in a CRPC-NEPC context, and PSMA2 loss re-sensitized enzalutamide-resistant cells. That said, the toxic interaction between ADT and bortezomib both in our mouse data and clinically^59^ strongly argues for next-generation proteasome-targeted agents with distinct subtype selectivity and broader therapeutic index.

Although bortezomib’s systemic toxicity limits its clinical utility, this should not discourage development of next-generation agents that selectively target PSMA2 or PSMA2-enriched proteasome subtypes. Subunit-selective proteasome inhibition has already been achieved. For example, LMP7-specific inhibitors with reduced toxicity^60^ and β-selective strategies^55^. These advances, together with emerging small-molecule and PROTAC-based approaches that modulate individual proteasome subunits^61^, demonstrate that precision targeting of PSMA2-containing proteasome assemblies is feasible and compatible with an improved therapeutic window. Overall, this work identifies PSMA2 as a central mechanistic and therapeutic node at the exact bottleneck between AR-driven and NEPC-driven disease. Targeting PSMA2 or the PSMA2-dependent proteasome subtype represents a rational strategy to intercept prostate cancer evolution at its most clinically consequential transition point, before AR-indifference becomes irreversible.

This work identifies PSMA2 as a treatment-induced driver of the conversion of AR-low to NEPC state. By enabling residual-androgen hypersensitization and simultaneously activating stemness/NEPC transcriptional programs, PSMA2 unifies two cardinal features of ADT resistance within a single mechanistic hub. Because PSMA2 induction occurs upstream of irreversible AR-indifference, PSMA2 expression or PSMA2-regulated proteasome subtype composition could serve as an early biomarker of impending lineage transition. Importantly, these findings point to PSMA2-dependent proteasome specialization as an actionable vulnerability for rational interception of tNEPC evolution in patients.

## ACKNOWLEDGMENT & FUNDING

The work was supported by NIH grants (T32-GM008396) and IMSD (R25 GM056901) Program to NP. Additional support was provided by MU start-up fund to CYC.

## AUTHORS CONTRIBUTION

NP: QPCR, Western blotting, preparation of manuscript and figures, AB: QPCR assays and Western Blot, GPV: FACS and co-immunoprecipitation experiments, RO: MTT assays and Western Blot, MH: Mouse colony maintenance and treatments, JM: Mouse colony maintenance and tumor harvesting, BD: sample processing for next generation sequencing. CYC: designed the study and wrote the manuscript.

## COMPETING INTERESTS

All authors declare no conflicts.

## MATERIALS AND METHODS

### Cell lines maintenance

All cell lines were obtained from ATCC (Manassas, VA, USA). LNCAP cell line was cultured in RMPI (Gibco #11875101) supplemented with 10% Fetal Bovine Serum (FBS). LASCPC-01 and NCI-H660 cells were cultured in custom RMPI (ATCC #30-2001) supplemented with 5% FBS, 10 nM hydrocortisone, 10 nM β-estradiol, 1x Insulin-Transferrin-Selenium, 1x GlutaMAX. All cells were maintained at 37°C with 5% CO2.

### Cell proliferation and drug sensitivity assays

#### MTT assays

Cell viability was performed using the MTT (3-(4,5-dimethylthiazol-2-yl)-2,5-diphenyl-2H-tetrazolium bromide) kit (Sigma #CT02) following manufacturer recommendation. In brief, 5×10^4^ LNCAP or NCI-H660 or LASCPC-01 cells were seeded in a 96-well plate and cultured for 24hrs, followed by treatment with Enzalutamide (1µm) and/or R1881(0.05-1nM), and/or Bortezomib (50-200nM) (LC LABS #B-1408) for 48hrs. MTT (5mg/ml) was added to the cultures, which were incubated for 4hrs. 200ul Acidic isopropanol was prepared as a 1.5% (v/v) solution of hydrochloric acid in isopropanol. Absorbance was read at 490nm on a microplate reader (BioTek Epoch).

#### Trypan Blue Assay

Following treatment (see above), cells were collected and centrifuged at 450g for 8 min, then resuspended in 1ml of culture media. 1-part cells were mixed with 1-part 0.4% trypan blue and incubated at room temperature for 3 mins. Viable cells were measured using Thermofisher automated cell counter.

### Transfection with short hairpin RNA interference or protein overexpression viral particles

Cells were seeded in a six-well plate at 5×10^5^ cells per well for 24hrs followed and transfected (Dharmafect 2) with shPSMA2 or with non-specific control shRNA (Horizon Dharmacon, PSMA2 lenti shRNA Transfection Starter kit # RHS11851-EG5683) or with Precision LentiORF PSMA2 viral particle starter kit (Horizon # OHS5900-202626736). Transfection and protein knockdown or overexpression efficiency was validated by immunoblot and QPCR analysis and Western blotting.

### Quantitative PCR analyses

Total RNA was extracted using an RNAeasy miniprep kit (Qiagen #74104) according to the manufacturer’s instructions. cDNA synthesis was performed using an Applied Biosystem High-Capacity RT kit (#4368814). Quantitative PCR was performed using Applied Biosystem SYBR Green PCR Kit. as per manufacturer’s instruction on BioRad CFX96™ System in 96-well plates in 3–6 repeats. A two-step thermal cycling protocol, i.e., 95 °C for 2 min followed by 50 cycles at 95 °C for 10 s and 56 °C for 60 s, was used. Fold change was calculated using the 2^−ΔΔCt^ method.

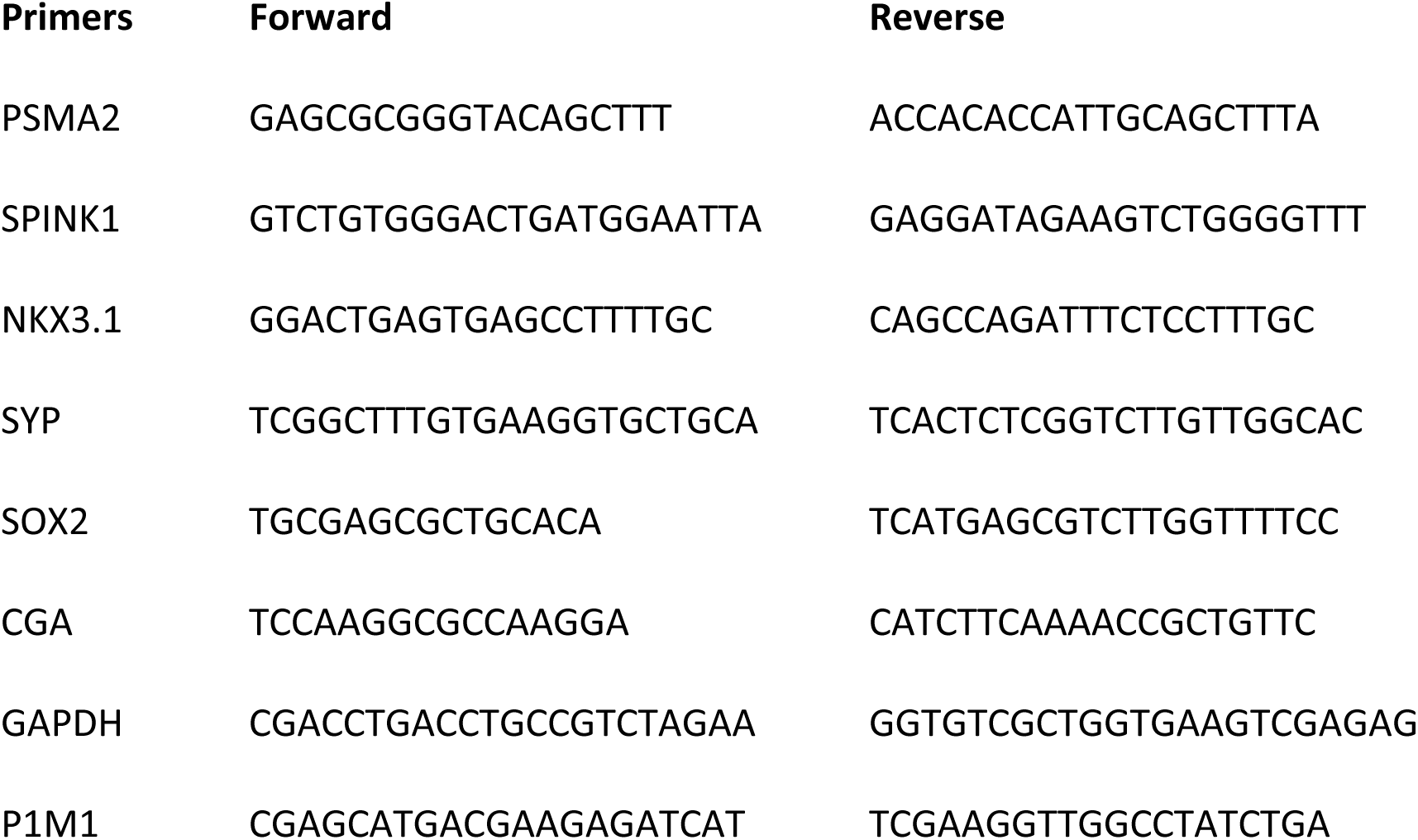

### Western blotting

Cells were lysed using Pierce RIPA Lysis and Extraction Buffer (ThermoFisher Scientific #89900) containing 1 × Halt™ Protease Inhibitor Cocktail (Thermofisher #78425). Blots were stained with antibodies against PSMA2 (Origene#CF505474), HSP90 (ThermoFisher Scientific # MA110372), PSA/KLK3 (Proteintech#60338-1), AR (Invitrogen # MA5-13426), and GAPDH (1:1000, DSHB-hGAPDH-2G7) or b-tubulin (1:1000, DSHB-E7). Secondary horseradish peroxidase (HRP) antibodies were obtained from Invitrogen. Pierce ECL Chemiluminescence kit (ThermoFisher Scientific #32106) and the ChemiDoc Imaging System (Bio-Rad) were used to detect protein bands.

### Immunohistochemistry

Cells were seeded at on Poly-L-lysine treated slide chambers followed by 48hr treatment. After treatment, cells were washed 3 times with cold PBS and fixed in 4% paraformaldehyde for 20 min. Cells were washed 3 times with cold PBS and permeabilized by incubation with cold 100% ethanol for 15 minutes at −20°C followed by washing with PBS. Cells were subsequently blocked for 1 h in PBS containing 0.01% Tween and 2% BSA and incubated with anti-synaptophysin (1:200, diluted in PBS containing 0.01% Triton tween and 2% BSA) for 1 hr at 37 °C. After three washes in PBS, cells were incubated for 15 min at 37 °C with anti-mouse Alexa Fluor 647 (1:400) secondary antibody, washed five times with PBS before mounting with VECTASHIELD mounting solution, and imaged on a Leica SP8 confocal microscope.

### Immunoprecipitation

Cells were lysed using Pierce RIPA Lysis and Extraction Buffer (ThermoFisher Scientific #89900) containing 1 × Halt™ Protease Inhibitor Cocktail (Thermofisher #78425). A total of 4mg protein was used. 100 μL of PBS washed A/G bead slurry (pierce Protein A/G agarose, Cat# 20421, Thermo Scientific) was used for 1mg/mL cell lysate. Cleared protein extracts were pre-incubated with AG beads for 1h at 4°C and incubated overnight at 4°C on a rotator with anti-AR primary antibodies (Invitrogen #MA5-13426, 1:1000). Beads were precipitated on a centrifuge (1,000g for 2 min at 4°C) and washed three times with wash buffer (10mM Tris-pH 7.4, 1mM EGTA, 150mM NaCl, 1%Triton X-100, with protease inhibitor cocktail). Binding proteins were eluted from the beads by heating or boiling samples in loading buffer with denaturant SDS and processes for SDSPAGE and Western blotting. Blots were stained with antibodies against AR (Invitrogen #MA5-13426, 1:1000), HSP90 (Invitrogen #MA110372, 1:500), GAPDH (DSHB #2G7, 1:1000), and anti β-actin (1/1,000; Sigma, cat# A5441).

### Fluorescence activated cell sorting

Cells were seeded for 24 at in a 6-well plate at 5x 10^5^ per well for 24 h, followed by treatment with Enzalutamide (1μm) and/or R1881(0.05-1nM) and/or Bortezomib (50-200nM) for 48hrs. After treatment, cells were stained for EDU, using Click-iT™ Plus EdU Alexa Fluor™ 647 Flow Cytometry Assay Kit (Molecular Probes C10634) following manufacturer protocol. Cells were analyzed using a FACS flow cytometer (BD Biosciences). Data analysis was performed using the FlowJo software (Tree Star; https://www.flowjo.com/).

### Mouse studies

Early tumor growth kinetics following PSMA2 knockdown in vivo. 10^5^ LASCPC-01 or LASCPC-01 PSMA2-KD cells were inoculated subcutaneously into the flanks of nude Foxn^1u^ (nu/nu) mice. Tumor volume was established by caliper measurement using the formula V = 0.5 x height x width^2^.

Kaplan-Meier survival studies for male Transgenic Adenocarcinoma of Mouse Prostate (TRAMP) mice (hybrid FvB x C57BL/6-Tg(TRAMP)8247Ng/J, JAX). Tumor positive 25-28 weeks old mice were randomized at disease onset and assigned to Untreated, Bortezomib (1 mg/kg, weekly), or Enzalutamide (3 mg/kg, provided *ad libitum* in drinking water and refreshed every 3-4 days) plus Bortezomib. Total treatment duration was 2 weeks, with mice followed until moribund. N = 6 per group. Statistical significance was assessed by log-rank (Mantel-Cox) test.

### Analysis of PSMA2 expression in prostate cancer patients

The Gene Expression Profiling Interactive Analysis (GEPIA2), http://gepia2.cancer-pku.cn/#analysis (Tang et al., 2019) was applied to compare PSMA2 mRNA levels in Prostate cancer tissue samples versus normal tissues using prostate cancer (PRAD) Cancer Genome Atlas and The Genotype-Tissue Expression (GTEx) with a threshold of p-value < .05 and Log2 fold change> 0.1. Survival analysis and expression NEPC markers (*MYCN, SPINK1, ONECUTE, REST, EZH2, FOXA2, AURKA, AKT1, LIN28B* and *SOX2*) in prostate cancer patients with PSMA2 amplification versus unaltered PSMA2 was performed on cBioPortal (http://cbioportal.org) ^52,53^.

## Notes

### Competing Interest Statement

The authors have declared no competing interest.

## REFERENCE

1 Vlachostergios, P. J. & Papandreou, C. N. Targeting neuroendocrine prostate cancer: molecular and clinical perspectives. Front Oncol 5, 6 (2015). 10.3389/fonc.2015.00006

2 Wang, H. T. et al. Neuroendocrine Prostate Cancer (NEPC) progressing from conventional prostatic adenocarcinoma: factors associated with time to development of NEPC and survival from NEPC diagnosis-a systematic review and pooled analysis. J Clin Oncol 32, 3383–3390 (2014). 10.1200/JCO.2013.54.3553

3 Beltran, H. et al. Challenges in recognizing treatment-related neuroendocrine prostate cancer. J Clin Oncol 30, e386–389 (2012). 10.1200/JCO.2011.41.5166

4 Bluemn, E. G. et al. Androgen Receptor Pathway-Independent Prostate Cancer Is Sustained through FGF Signaling. Cancer Cell 32, 474–489 e476 (2017). 10.1016/j.ccell.2017.09.003

5 Aggarwal, R. et al. Clinical and Genomic Characterization of Treatment-Emergent Small-Cell Neuroendocrine Prostate Cancer: A Multi-institutional Prospective Study. J Clin Oncol 36, 2492–2503 (2018). 10.1200/JCO.2017.77.6880

6 Lundberg, A. et al. The Genomic and Epigenomic Landscape of Double-Negative Metastatic Prostate Cancer. Cancer Res 83, 2763–2774 (2023). 10.1158/0008-5472.CAN-23-0593

7 Labrecque, M. P. et al. Molecular profiling stratifies diverse phenotypes of treatment-refractory metastatic castration-resistant prostate cancer. J Clin Invest 129, 4492–4505 (2019). 10.1172/JCI128212

8 Beltran, H. et al. Divergent clonal evolution of castration-resistant neuroendocrine prostate cancer. Nat Med 22, 298–305 (2016). 10.1038/nm.4045

9 Alumkal, J. J. et al. Transcriptional profiling identifies an androgen receptor activity-low, stemness program associated with enzalutamide resistance. Proc Natl Acad Sci U S A 117, 12315–12323 (2020). 10.1073/pnas.1922207117

10 Pratt, W. B. & Toft, D. O. Steroid receptor interactions with heat shock protein and immunophilin chaperones. Endocr Rev 18, 306–360 (1997). 10.1210/edrv.18.3.0303

11 Chen, Y., Sawyers, C. L. & Scher, H. I. Targeting the androgen receptor pathway in prostate cancer. Curr Opin Pharmacol 8, 440–448 (2008). 10.1016/j.coph.2008.07.005

12 Veldscholte, J., Berrevoets, C. A., Brinkmann, A. O., Grootegoed, J. A. & Mulder, E. Anti-androgens and the mutated androgen receptor of LNCaP cells: differential effects on binding affinity, heat-shock protein interaction, and transcription activation. Biochemistry 31, 2393–2399 (1992). 10.1021/bi00123a026

13 Kuil, C. W., Berrevoets, C. A. & Mulder, E. Ligand-induced conformational alterations of the androgen receptor analyzed by limited trypsinization. Studies on the mechanism of antiandrogen action. J Biol Chem 270, 27569–27576 (1995). 10.1074/jbc.270.46.27569

14 Georget, V., Terouanne, B., Nicolas, J. C. & Sultan, C. Mechanism of antiandrogen action: key role of hsp90 in conformational change and transcriptional activity of the androgen receptor. Biochemistry 41, 11824–11831 (2002). 10.1021/bi0259150

15 Trepel, J., Mollapour, M., Giaccone, G. & Neckers, L. Targeting the dynamic HSP90 complex in cancer. Nat Rev Cancer 10, 537–549 (2010). 10.1038/nrc2887

16 Beltran, H. et al. The Role of Lineage Plasticity in Prostate Cancer Therapy Resistance. Clin Cancer Res 25, 6916–6924 (2019). 10.1158/1078-0432.CCR-19-1423

17 Mu, P. et al. SOX2 promotes lineage plasticity and antiandrogen resistance in TP53- and RB1-deficient prostate cancer. Science 355, 84–88 (2017). 10.1126/science.aah4307

18 Romero, R. et al. The neuroendocrine transition in prostate cancer is dynamic and dependent on ASCL1. Nat Cancer 5, 1641–1659 (2024). 10.1038/s43018-024-00838-6

19 Beltran, H. et al. Aggressive variants of castration-resistant prostate cancer. Clin Cancer Res 20, 2846–2850 (2014). 10.1158/1078-0432.CCR-13-3309

20 Linder, S., van der Poel, H. G., Bergman, A. M., Zwart, W. & Prekovic, S. Enzalutamide therapy for advanced prostate cancer: efficacy, resistance and beyond. Endocr Relat Cancer 26, R31–R52 (2018). 10.1530/ERC-18-0289

21 Wang, G., Zhao, D., Spring, D. J. & DePinho, R. A. Genetics and biology of prostate cancer. Genes Dev 32, 1105–1140 (2018). 10.1101/gad.315739.118

22 Berger, A. et al. N-Myc-mediated epigenetic reprogramming drives lineage plasticity in advanced prostate cancer. J Clin Invest 129, 3924–3940 (2019). 10.1172/JCI127961

23 Tiwari, R. et al. Androgen deprivation upregulates SPINK1 expression and potentiates cellular plasticity in prostate cancer. Nat Commun 11, 384 (2020). 10.1038/s41467-019-14184-0

24 Yamada, Y. & Beltran, H. Clinical and Biological Features of Neuroendocrine Prostate Cancer. Curr Oncol Rep 23, 15 (2021). 10.1007/s11912-020-01003-9

25 Glickman, M. H. & Ciechanover, A. The ubiquitin-proteasome proteolytic pathway: destruction for the sake of construction. Physiol Rev 82, 373–428 (2002). 10.1152/physrev.00027.2001

26 Ugai, S. et al. Purification and characterization of the 26S proteasome complex catalyzing ATP-dependent breakdown of ubiquitin-ligated proteins from rat liver. J Biochem 113, 754–768 (1993). 10.1093/oxfordjournals.jbchem.a124116

27 Arlt, A. et al. Increased proteasome subunit protein expression and proteasome activity in colon cancer relate to an enhanced activation of nuclear factor E2-related factor 2 (Nrf2). Oncogene 28, 3983–3996 (2009). 10.1038/onc.2009.264

28 Tamura, T. et al. Molecular cloning and sequence analysis of cDNAs for five major subunits of human proteasomes (multi-catalytic proteinase complexes). Biochim Biophys Acta 1089, 95–102 (1991). 10.1016/0167-4781(91)90090-9

29 Coux, O., Tanaka, K. & Goldberg, A. L. Structure and functions of the 20S and 26S proteasomes. Annu Rev Biochem 65, 801–847 (1996). 10.1146/annurev.bi.65.070196.004101

30 Ravid, T. & Hochstrasser, M. Diversity of degradation signals in the ubiquitin-proteasome system. Nat Rev Mol Cell Biol 9, 679–690 (2008). 10.1038/nrm2468

31 Sun, P. et al. Bufalin derivative BF211 inhibits proteasome activity in human lung cancer cells in vitro by inhibiting beta1 subunit expression and disrupting proteasome assembly. Acta Pharmacol Sin 37, 908–918 (2016). 10.1038/aps.2016.30

32 Chiao, C. C. et al. Prognostic and Genomic Analysis of Proteasome 20S Subunit Alpha (PSMA) Family Members in Breast Cancer. Diagnostics (Basel) 11 (2021). 10.3390/diagnostics11122220

33 Sandhu, H. S. et al. Dynamic plasticity of prostate cancer intermediate cells during androgen receptor-targeted therapy. Cell Rep 40, 111123 (2022). 10.1016/j.celrep.2022.111123

34 Lee, J. K. et al. N-Myc Drives Neuroendocrine Prostate Cancer Initiated from Human Prostate Epithelial Cells. Cancer Cell 29, 536–547 (2016). 10.1016/j.ccell.2016.03.001

35 Cerasuolo, M. et al. Neuroendocrine Transdifferentiation in Human Prostate Cancer Cells: An Integrated Approach. Cancer Res 75, 2975–2986 (2015). 10.1158/0008-5472.CAN-14-3830

36 Shen, R. et al. Transdifferentiation of cultured human prostate cancer cells to a neuroendocrine cell phenotype in a hormone-depleted medium. Urol Oncol 3, 67–75 (1997). 10.1016/s1078-1439(97)00039-2

37 Dankert, J. T. et al. The deregulation of miR-17/CCND1 axis during neuroendocrine transdifferentiation of LNCaP prostate cancer cells. PLoS One 13, e0200472 (2018). 10.1371/journal.pone.0200472

38 Chiaverotti, T. et al. Dissociation of epithelial and neuroendocrine carcinoma lineages in the transgenic adenocarcinoma of mouse prostate model of prostate cancer. Am J Pathol 172, 236–246 (2008). 10.2353/ajpath.2008.070602

39 Gingrich, J. R. et al. Androgen-independent prostate cancer progression in the TRAMP model. Cancer Res 57, 4687–4691 (1997).

40 Gingrich, J. R. et al. Metastatic prostate cancer in a transgenic mouse. Cancer Res 56, 4096–4102 (1996).

41 Kazmierczak, R. A. et al. Evaluations of CRC2631 toxicity, tumor colonization, and genetic stability in the TRAMP prostate cancer model. Oncotarget 11, 3943–3958 (2020). 10.18632/oncotarget.27769

42 Franko, A. et al. Characterization of Hormone-Dependent Pathways in Six Human Prostate-Cancer Cell Lines: A Gene-Expression Study. Genes (Basel) 11 (2020). 10.3390/genes11101174

43 Liu, Y. N. et al. Leukemia Inhibitory Factor Promotes Castration-resistant Prostate Cancer and Neuroendocrine Differentiation by Activated ZBTB46. Clin Cancer Res 25, 4128–4140 (2019). 10.1158/1078-0432.CCR-18-3239

44 Zegarra-Moro, O. L., Schmidt, L. J., Huang, H. & Tindall, D. J. Disruption of androgen receptor function inhibits proliferation of androgen-refractory prostate cancer cells. Cancer Res 62, 1008–1013 (2002).

45 Mohler, J. L. et al. The androgen axis in recurrent prostate cancer. Clin Cancer Res 10, 440–448 (2004). 10.1158/1078-0432.ccr-1146-03

46 Stanbrough, M. et al. Increased expression of genes converting adrenal androgens to testosterone in androgen-independent prostate cancer. Cancer Res 66, 2815–2825 (2006). 10.1158/0008-5472.CAN-05-4000

47 Montgomery, R. B. et al. Maintenance of intratumoral androgens in metastatic prostate cancer: a mechanism for castration-resistant tumor growth. Cancer Res 68, 4447–4454 (2008). 10.1158/0008-5472.CAN-08-0249

48 Chen, C. D. et al. Molecular determinants of resistance to antiandrogen therapy. Nat Med 10, 33–39 (2004). 10.1038/nm972

49 Xie, Q. & Wang, Z. A. Transcriptional regulation of the Nkx3.1 gene in prostate luminal stem cell specification and cancer initiation via its 3’ genomic region. J Biol Chem 292, 13521–13530 (2017). 10.1074/jbc.M117.788315

50 Carver, B. S. Defining and Targeting the Oncogenic Drivers of Neuroendocrine Prostate Cancer. Cancer Cell 29, 431–432 (2016). 10.1016/j.ccell.2016.03.023

51 Zou, M. et al. Transdifferentiation as a Mechanism of Treatment Resistance in a Mouse Model of Castration-Resistant Prostate Cancer. Cancer Discov 7, 736–749 (2017). 10.1158/2159-8290.CD-16-1174

52 Cerami, E. et al. The cBio cancer genomics portal: an open platform for exploring multidimensional cancer genomics data. Cancer Discov 2, 401–404 (2012). 10.1158/2159-8290.CD-12-0095

53 Gao, J. et al. Integrative analysis of complex cancer genomics and clinical profiles using the cBioPortal. Sci Signal 6, pl1 (2013). 10.1126/scisignal.2004088

54 Adams, J. The proteasome: a suitable antineoplastic target. Nat Rev Cancer 4, 349–360 (2004). 10.1038/nrc1361

55 Kisselev, A. F. & Goldberg, A. L. Proteasome inhibitors: from research tools to drug candidates. Chem Biol 8, 739–758 (2001). 10.1016/s1074-5521(01)00056-4

56 Bross, P. F. et al. Approval summary for bortezomib for injection in the treatment of multiple myeloma. Clin Cancer Res 10, 3954–3964 (2004). 10.1158/1078-0432.CCR-03-0781

57 Sruthi, K. K. & Ummanni, R. Valosin-Containing Protein (VCP/p97) Mediates Neuroendocrine Differentiation in Prostate Cancer Cells Through Pim1 Signaling Inducing Autophagy. Prostate 85, 932–946 (2025). 10.1002/pros.24900

58 Vlachostergios, P. J. & Papandreou, C. N. Neuroendocrine differentiation and the ubiquitin-proteasome system in cancer: Partners or enemies? World J Exp Med 1, 7–9 (2011). 10.5493/wjem.v1.i1.7

59 Kraft, A. S. et al. Combination therapy of recurrent prostate cancer with the proteasome inhibitor bortezomib plus hormone blockade. Cancer Biol Ther 12, 119–124 (2011). 10.4161/cbt.12.2.15723

60 Muchamuel, T. et al. A selective inhibitor of the immunoproteasome subunit LMP7 blocks cytokine production and attenuates progression of experimental arthritis. Nat Med 15, 781–787 (2009). 10.1038/nm.1978

61 Leestemaker, Y. et al. Proteasome Activation by Small Molecules. Cell Chem Biol 24, 725–736 e727 (2017). 10.1016/j.chembiol.2017.05.010

